# Modelling Emergence of *Wolbachia* Toxin-Antidote Protein Functions with an Evolutionary Algorithm

**DOI:** 10.1101/2023.03.23.533954

**Authors:** John Beckmann, Joe Gillespie, Daniel Tauritz

## Abstract

Evolutionary algorithms (EAs) simulate Darwinian evolution and adeptly mimic natural evolution. Most EA applications in biology encode high levels of abstraction in top-down ecological population models. In contrast, our research merges protein alignment algorithms from bioinformatics into codon based EAs that simulate molecular protein string evolution from the bottom up. We apply our EA to reconcile a problem in the field of *Wolbachia* induced cytoplasmic incompatibility (CI). *Wolbachia* is a microbial endosymbiont that lives inside insect cells. CI is conditional insect sterility that operates as a toxin antidote (TA) system. Although, CI exhibits complex phenotypes not fully explained under a single discrete model. We instantiate in-silico genes that control CI, CI factors (*cifs*), as strings within the EA chromosome. We monitor the evolution of their enzymatic activity, binding, and cellular localization by applying selective pressure on their primary amino acid strings. Our model helps rationalize why two distinct mechanisms of CI induction might coexist in nature. We find that nuclear localization signals (NLS) and Type IV secretion system signals (T4SS) are of low complexity and evolve fast, whereas binding interactions have intermediate complexity, and enzymatic activity is the most complex. Our model predicts that as ancestral TA systems evolve into eukaryotic CI systems, the placement of NLS or T4SS signals can stochastically vary, imparting effects that might impact CI induction mechanics. Our model highlights how preconditions, genetic diversity, and sequence length can bias evolution of *cifs* towards one mechanism or another.

## Introduction

TA systems typically involve two linked genes encoding a toxin and antidote.^1^ They skew Mendelian inheritance in their favor by addicting organisms to the presence of an antidote and killing offspring that don’t inherit the TA module, via the toxin. Thus, they ensure inheritance in the next generation by post segregational killing. Ancestrally, TA systems might have arisen as selfish systems linked to the replication of prokaryotic plasmids.^2^ How TA systems evolve is a chicken-egg paradox: a lone toxin is detrimental to host fitness and an antidote without a linked cognate toxin could be beneficial, neutral, or detrimental, dependent on context. Prior models predict that TA systems evolve at lower levels of selection on plasmids; in contexts of genomic conflict or in situations where antidotes have beneficial functions in addition to toxin rescue.^2^

*Wolbachia* are bacteria that live inside insects.^3-5^ *Wolbachia* have the capability to sterilize mosquitos in a phenotype called CI.^6-10^ CI is a unique biological instantiation of a TA system. The CI phenotype is useful ^8,11,12^. CI is currently applied as a biocontrol mechanism preventing the transmission of mosquito borne diseases across the world and on multiple continents in various applications. Mosquitos infected with *Wolbachia* exhibit reduced ability to transmit flaviviruses like Dengue and Zika ^13,14^. Ongoing attempts use the selective pressure of *cifs* to spread beneficial (probiotic) *Wolbachia* infections into wild mosquito populations to limit disease ^15^. At the molecular level, the beneficial spread of *Wolbachia* is linked to the function of *cif* TA genes.

Much of the evolutionary dynamics of CI has been well described at the population level. CI is common because it increases equilibrium frequencies and infection persistence, thereby increasing the chances of *Wolbachia* being transferred to new species hosts.^16^ Yet in the insect, selection does not act to preserve or increase CI rates.^17^ Importantly, evolutionary dynamics and selective pressures operating at the lowest molecular level and at the moment CI emerged in evolutionary history have never been described.

The genes that control this conditional sterility are two linked genes dubbed *cifs* that form complex TA systems.^18-20^ Uniquely, this TA system is both bacterial and eukaryotic because it is encoded within intracellular bacterial endosymbionts yet expresses extended phenotypes impacting the eukaryotic insect host. *Cifs* are uniquely positioned in that their evolutionary origin necessitates a functional jump from bacteria to eukaryotes. The *cif* TA system encodes a sperm delivered embryo killer toxin and a cognate rescuing antidote. If the insect host loses *Wolbachia*, remaining toxin sterilizes males, and these populations don’t reproduce. Therefore, female insects keep *Wolbachia* because the antidotes are useful in the presence of toxins encountered in male sperm.

Importantly, purifying selection does preserve the *cif* antidotes;^21-23^ and on lower levels in the context of genomic conflict, selection can act to assemble the biochemical domains of toxin antidote systems.^2^ Though once assembled, selection on the insect level does not act to preserve the bacterial toxins which tend to pseudogenize and/or are replaced by subsequent invading *cif* systems.^24,25^

While molecular details on CI function are emerging, one problem is that rules governing induction of sterility via the *Wolbachia* TA system are debated. In general, the system behaves as a classical TA module, meaning one gene named *cifB* is inducer and its cognate partner *cifA* acts as antidote.^20^ However, there remains unresolved nuance in the mechanism. Currently all data support the hypothesis that the first operon gene, *cifA*, is antidote.^19,26^ However, induction of CI and the exact source of the toxicity appears more, or less, complex in various models. The two main models each have empirical evidence to support them. These models are the TA model^20,27,28^ and the 2×1 model.^29,30^ The TA model is more parsimonious and significant evidence supports it in fruit flies, mosquitos, yeast models, and structural studies^19,31-36^. In contrast, the 2×1 model posits that a single gene acts as rescue factor, but induction of sterility requires both *cifA* and *cifB* genes.^29,37^

Our hypothesis is that both models coexist in nature as alternate variations of the broader TA theme. These variations might arise as CI evolves from a simple prokaryotic TA module into a eukaryotic CI system (see **Fig 1A** and **B**). To explain, induction of sperm sterility in a eukaryote via a prokaryotic TA module necessitates the evolution of additional functions beyond toxin and antidote. In support of this hypothesis, prior models predicted that beneficial functions in addition to antidote functionality are prerequisites for TA module emergence.^2^ In our case, *Wolbachia* must first secrete the toxin out one of its Sec-independent secretion systems; for the remainder of this study, we implicate the Rickettsiales *vir* homolog (*rvh*) type IV secretion system (T4SS) for CI protein secretion. T4SS substrates require a signal sequence, usually found at the C-terminus. Once secreted, the toxin must localize into the nucleus via a nuclear localization signal (NLS). There is a likely possibility that binding of *cifA* to *cifB* occurs prior to secretion and thus one protein might drag the other through a given secretion system. Under this hypothesis, it is possible that T4SS and NLS sequences could evolve in either antidote or toxin genes in different insect hosts. If the *cifA* antidote acquires an NLS and T4SS signal but *cifB* has neither, this leads to additional complexity in the system necessitating cooperative induction of sterility by *cifA* and *cifB* (hence a 2×1). While most empirical work evidences a strict TA in four known orthologs (cid^*w*Pip^, cid^*w*Ha^, cin^*w*No^, and cin^Ott^), there is indication of 2×1 in two systems (cid^*w*Mel^ and cin^*w*Pip^). Our research did not focus on determining if one model was correct at the complete expense of the other, but rather seeks to understand evolutionary pressures and selective mechanisms that might bias evolution of one model over another. Understanding the precise molecular mechanisms underlying the *cif* TA system and its evolution contributes information to “fine-tune” *Wolbachia* based biocontrol. Once we have perfect knowledge for how the *cif* TA sterility is induced, we can design the most efficient and parsimonious transgene insertions to reconstruct sterility in transgenic mosquitos as a biotechnological tool (i.e., with 2 genes or 1).

**Figure 1:**
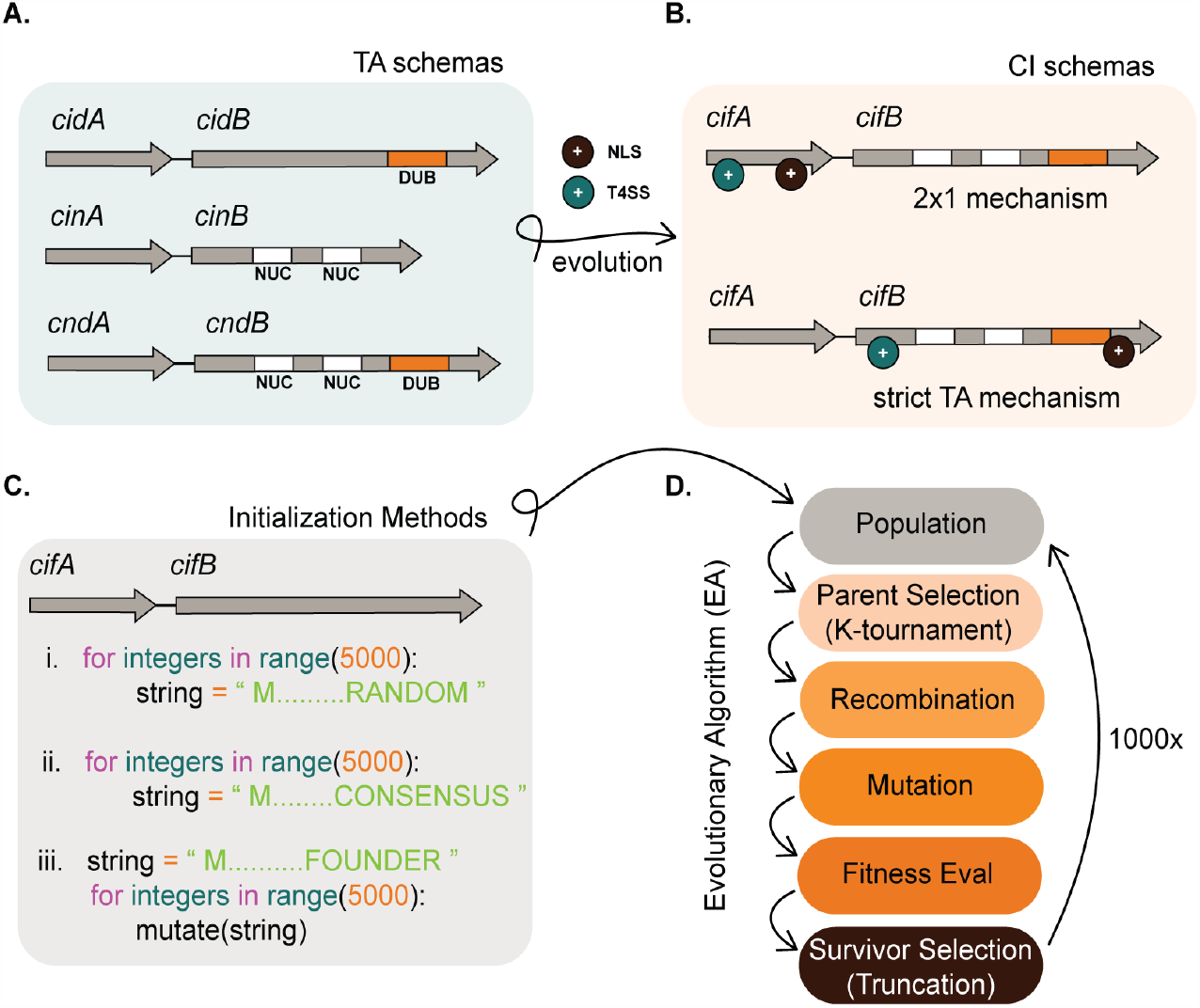
Background infographic. **A**. Schemas of *Wolbachia* TA modules in a more ancestral prokaryotic form. To evolve into CI systems, ancestral TA modules must add more complex features including an NLS (+ black circle) and a T4SS (+ cyan circle). **B**. CI system schemas might evolve into two descriptive models which include the 2×1 and strict TA model. The location where NLS or T4SS features evolve could impact the mechanistic induction of CI. A CI schema where both NLS and T4SS features co-occur in *cifA* alone is predicted to require both *cifA* and *cifB* for induction. In contrast, if these features co-occur in *cifB*, then *cifB* would be sufficient for induction of CI and behave as a strict TA module. **C**. In-Silico simulation of this evolution requires an initial instantiation of a population of TA strings. Our experiments tested three distinct methods of instantiation that include (i) instantiating random strings, (ii) instantiating semi-random strings comporting to conserved *cif* consensus sequence, and (iii) instantiating a single individual and deriving an entire population by mutagenesis of that founder. **D**. After instantiation of the population, it evolves under the selective pressure of a fitness function and follows discrete generations. Our algorithm selects parents by K-tournament and distributes these individuals into a mating pool. Offspring are generated by recombination of parents wherein two strings swap discrete sub-strings to create a new child. After recombination, child strings are mutated. Fitness of the TA is then evaluated, and survivors are selected based upon truncation survivor selection. In truncation, the population is sorted and the lowest fitness individuals that fall below a threshold are culled such that population numbers remain at the carrying capacity. The algorithm terminates after 1000x generations.

It was our goal to gain insights on the molecular evolution of CI by modelling CI’s emergence with an evolutionary algorithm. Using EAs to model natural evolution has been a productive application.^38-41^ Modelling gene drives in mosquitos with EAs and machine learning provided insights that predicted efficacy of actual biocontrol tools.^42,43^ However, biological evolution can be modelled by EAs at different ecological levels. Various abstractions and assumptions are made by any given model. EAs are typically top-down ecological models modelling populations of organisms. Top down EAs model gene flow of beneficial or deleterious traits. Within populations, each organism can be assigned a fitness value. Organisms and their genes can then mate, recombine, mutate, and die. These EA implementations tend towards Wright-Fisher models and often obey set rules. These models are useful for questions on evolutionary theory and adeptly model gene drives and selective sweeps, etc. However, the actual coded implementations are often abstract and difficult to translate into the evolution of amino acid sequence.

Popular bottom-up EA frameworks that modeled evolution upwards towards complexity are also abstract because they implemented computer assembly functions like “push” and “pop” as analogies of protein and metabolic pathways.^38,44^ These studies have demonstrated that in bottom-up simulations, simple functions can give rise to more complex functions (like add and multiply) through evolution; however, these are abstract analogies, not actual DNA code. There is a gap in implementations of bottom-up biological models. Bottom-up implementations could implement DNA code as the starting point and model how code changes. A bottom-up implementation should instantiate the lowest levels of selection on actual genes^45^ and test the lowest level of function which is protein translations of that code. The in-silico genes could be mutated and recombined as actual DNA molecules and fitness can determined by bioinformatic algorithms comparing string sequence similarity to proteins of known function. Our coded framework presented herein is novel in this respect.

EAs are perfect for studying protein string evolution because the search space of protein strings is vast (considering 20 possible amino acids and strings in lengths of thousands = 20^1000^ unique strings). Research implementing codon based EAs is in early stages.^46-49^ For example, a few studies tested machine learning guided mutations and used EAs to design novel antimicrobial peptides (AMPs).^46,48,49^ These researchers guided evolution from known AMP strings rather than evolving novel de-novo protein strings. These studies provide some support for the concept of using sequence similarity as a proxy for fitness. Here we use sequence similarity to *cif* consensus sequences as fitness proxy to process simulations orders of magnitude faster than possible with bioassay. To gain a better understanding of *cif* evolution, we encoded *cif* TA genes directly as chromosomes within an EA and observed the evolution of their strings (see **Fig 1C** and **D**).

## Materials and Methods

### EA Design

An overlapping generations (μ+λ) EA was coded in Python 3^1^ where population size (μ) was 5000 individuals and offspring (λ) was 100. Other variations were tested (see **Fig 2A**). Within the EA, code classes included an EA class (running the EA simulation functions, main methods, and data logging capacities), a TA class (housing the chromosome instantiations), and a main driver. The driver receives input from an editable JSON configuration file. All configuration files and outputs were saved and stored for reproducibility. The random seed is configurable for reproducibility.

**Figure 2:**
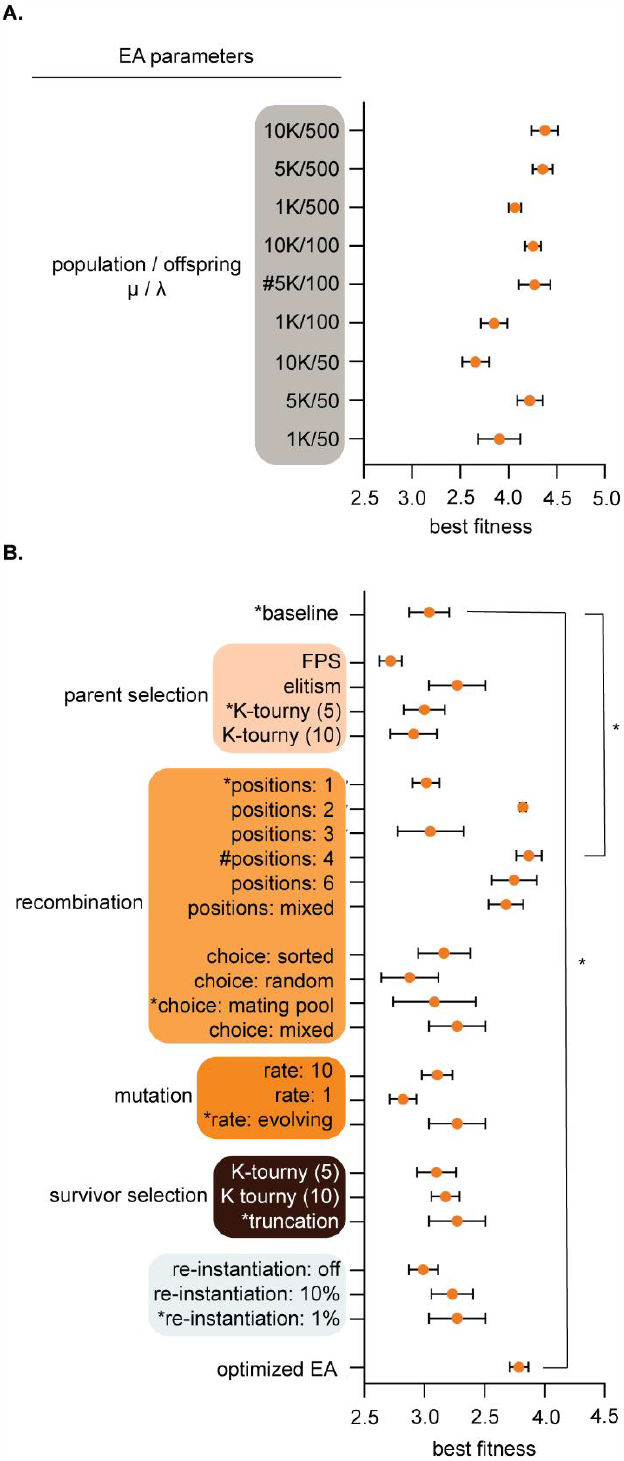
EA parameters were optimized for evolution and computational speed. **A**. Population parameters were pre-tested to configure population (μ) and offspring (λ) sizes for subsequent larger experiments. Parameters (Y-axis) and best individual fitness (X-axis) were logged after 100 generations of simulated evolution. We chose 5K/100 [see hashtag (#)] for population/offspring (μ/λ) because it yielded high fitness, diverse outcomes, and fast computation time. **B**. EA parameters were tuned by recording best fitness after populations [μ/ λ: 1000/100] were evolved for 100 generations. A baseline configuration (asterisks, *) was held constant while individual parameters were varied. Choosing the highest yielding fitness configuration for each parameter is shown at bottom as the “optimized EA”, though this was not necessarily *the best* because parameters exhibited interdependence. The optimized EA and baseline with 4-point crossover recombination evolved significantly better than baseline p<0.0001 by One-way ANOVA with Tukey post-hoc analysis. We used the baseline configuration with 4-point crossover recombination for subsequent experiments. Results show means and standard deviation from five trial runs after 100 generations.

### EA Class

EA algorithms are stochastic in nature. Evolutionary trajectories can proceed down different routes or converge. Thus, our main EA experiments consisted of 30 runs each (**Figs 3** and **4**). Main methods within the EA class included class resets (to reset logs and class variables after each run); population instantiations [including i), ii) and iii) see **Fig 1C**]; sorting functions that sorted TA populations based on fitness; parent selection methods (which finally used K-tournament of K = 5 after preliminary testing; see **Fig 2B**). K-tournament selection runs multiple fitness tournaments among a few individuals chosen randomly from the population. Winners of a tournament with the best fitness are sent to the mating pool array to be selected for recombination. In experiments recombination used 4-point crossover recombination, but we tested other modes (**Fig 2B**). Mutation methods utilized an algorithm that randomly locates DNA base pairs and flips to a random choice of A, T, G, or C. Mutation also encoded insertion and deletion functions with randomly sized indels. The mutation method evaluates fitness by calling the fitness evaluation method from the TA class (see below). After fitness evaluation, survivors were selected via truncation survivor selection. Other survivor selection regimes were tested (**Fig 2B**). Truncation sorts the population and culls the lowest fitness individuals in a number equivalent to the number of offspring added per generation. Thus, carrying capacity remains constant at μ = 5000. Data logging functions were encoded, for example, calculateAverageFitness(), which tallies an average TA fitness. A termination condition method was coded but not used in final experiments. Logs were recorded in output files and saved. We tracked 15 quantifiable observations: 1) highestTAFitness_HTF, 2) avgBindingFitness_ABF, 3) avgDUBFitness_ADF, 4) avgNucFitness_ANF, 5) avgTAfitness_ATF, 6) avgToxinLength_ATL, 7) avgToxinAALength_ATAL, 8) avgAntidoteLength_AAL, 9) avgAntidoteAALength_AAAL, 10) avgTAMutationRate_ATMR, 11) avgNLSSITELocation_NLSL, 12) avgTypeIVSITELocation_TYPL, 13) avgNLSFitness_ANLSF, 14) avgT4SSFitness_AT4F, 15) diversityIndex_DI. To briefly explain algorithmic terminology in **Fig 2**., *FPS* is fitness proportional parent selection which assigns mating probability as proportional to fitness; *Elitism* ranks parents on fitness and sends the most fit individuals into the mating pool, *K-tournament* is described above, *recombination* swaps DNA from two mated individuals at 1, 2, 3, 4, 6, or mixed points respectively; *mating choice sorted* sorts the mating pool and individuals mate with a partner closest to their fitness score, *mating choice random* allows individuals within the mating pool to randomly pick any other mate in the population; *mating choice mating pool* allows random choice of mates from within the mating pool only; *mating choice mixed* rolls a dice and chooses any method stochastically, *mutation rate* number indicates the number of dice rolls each individual child undergoes for chances to iteratively mutate the chromosome (the dice is an equal probability of 4 options to do nothing, bit flip, insert, or delete), *truncation* is described above, *re-instantiation* is a method to maintain diversity and it instantiates new TA modules from scratch and allows them to immigrate into the population at a set OKach generation.

**Figure 3:**
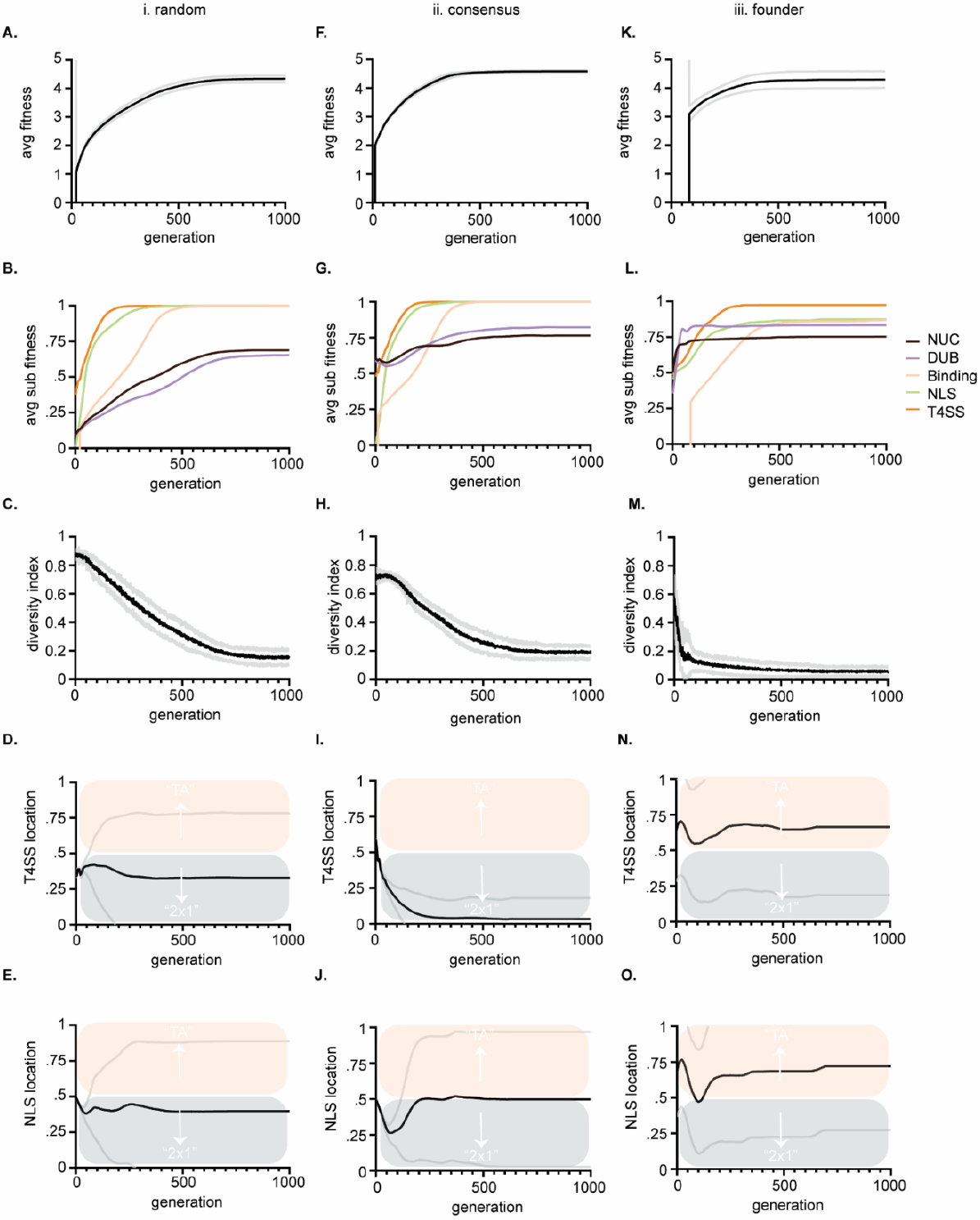
Output data from three large evolution experiments. Columns (i-iii) show results from three different instantiation methods described in **Fig 1**. Rows show average fitness of all TAs within a population versus generations (**A, F**, K). Average sub-feature fitness of all TAs within a population versus generations (**B, G, L**); Diversity index versus generations, where lower numbers indicate more similarity in string sequence and therefore loss of diversity (**C, H, M**); Average T4SS site location of the population versus generations (**D, I, N**); Average NLS site location of the population versus generations (**E, J, O**). Scoring for T4SS and NLS sites is as follows: a score of 0 indicates that the site evolved in the antidote gene (*cifA*) and a 1 indicates that the site evolved in the toxin gene (*cifB*); therefore a score of 0.5 means that half the population had the site in *cifA* and the other half had the site in *cifB*. Mean values are plotted with black or colored lines. Standard deviation is marked in grey lines. Bias above 0.5 indicates preferential evolution of TA and below 0.5, 2×1. **D, I**, and **N** are all significantly different from each other at termination, (p<0.05) by one-way ANOVA with Tukey post-hoc analysis. **E** and **O** were significantly different by the same.

**Figure 4:**
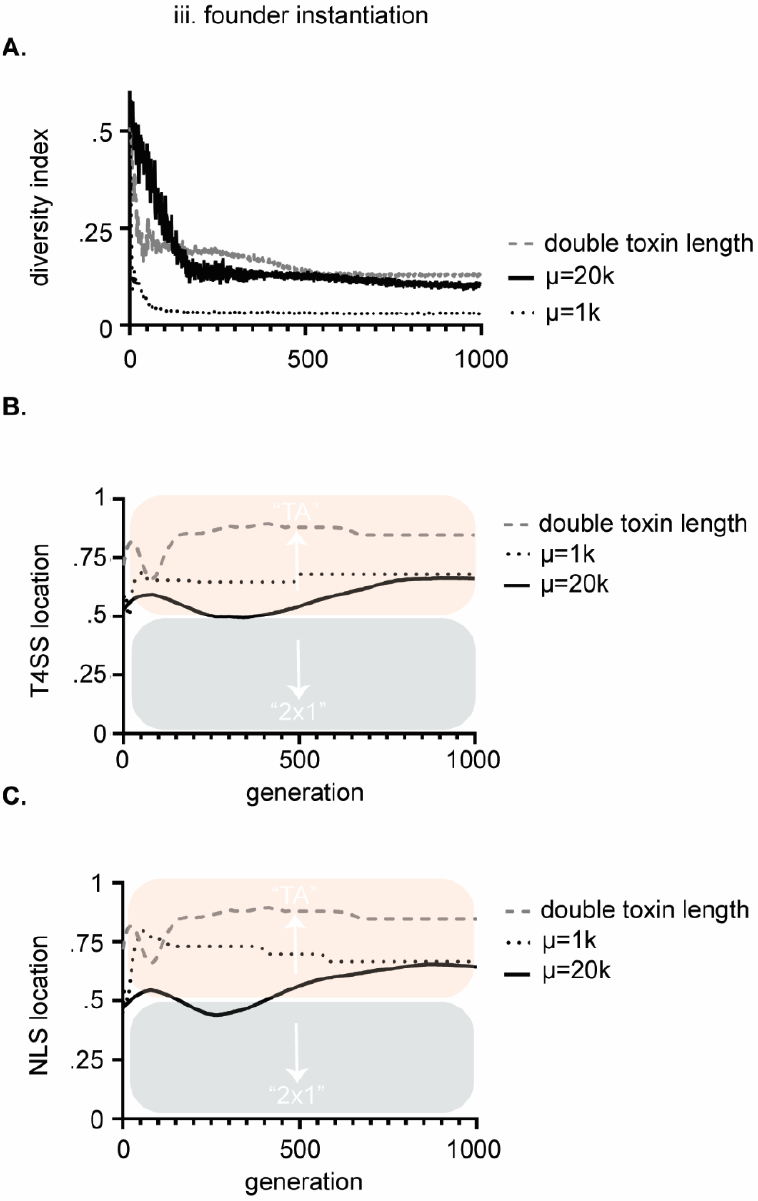
Parameters that control bias in the location of NLS and T4SS signals were tested. **A**. We were able to control starting diversity by changing population size (μ). Small populations of 1000 had significantly less diversity than populations of 20,000 by Mann-Whitney U test. In turn, these changes significantly altered the bias of the signal during the course of evolution, but not the final result (see dotted line vs solid line in **B**. and **C**.; T4SS location and NLS location respectively). The most significant terminal impact on bias of signal location was when we increased the average length of the toxins (see dashed line in **B** and **C**). **B**. and **C**. are T4SS and NLS sites is as above. Mean values are plotted with black or colored lines. In **B**. terminal conditions of doubling toxin length to µ=20k were significantly different for T4SS signal locations (p<0.05) by unpaired t-test with Welch’s correction. These experiments all were performed under the third [iii) founder] method of instantiation.

### TA Class

TA class individuals were instantiated with chromosomes encoding the string toxin and string antidote in DNA code. TAs additionally hold class variables including a nuclease score (measuring how well the toxin schema matches a known *cin* toxin consensus sequence) and a deubiquitylating (DUB) score (measuring how well the toxin schema matches a known *cid* toxin consensus sequence).^50^ They also hold a NLS score which is determined by presence or absence of a “KRAR” string ^51^ and a T4SS score determined by presence or absence of a “R-X(7)-R-X-R-X-R” string.^52^ All functional domains including nuclease domain, deubiquitylating domain, NLS, and T4SS signals are detected through a pairwise alignment algorithm and can be given partial scores if parts of the sequence are present. Pairwise alignment is built into the EA by importation of the Biopython^2^ module’s pairwise2 method. “Biopython is a set of freely available tools for biological computation written in Python by an international team of developers.” The pairwise2 method is called with a -1 gap penalty, a -0.1 gap extension penalty, and a false condition so that end gaps are not penalized. A binding score (measuring how well the pair bind each other) is determined by our own algorithm. This algorithm is based on a sliding window that slides two strings together in comparison to find and tally a score of the best matching residue configurations. Precisely 11 charged residues are known to underlie *cifA* and *cifB* binding ^35^. Therefore, if a sliding window detects an alignment of K with D, a score would be increased by 1 and the process continues. Repelling charges are penalized by -1. A total matched binding sequence shouldn’t exceed 11 binding residues in accordance with crystal structure data.^35^ Class methods within the TA class include standard “getters” and “setters” (i.e., setSchemata() which instantiates the toxin strings), a translation method that translates the DNA code into proteins, a coded number parser to facilitate binding evaluations with integers rather than strings (to speed up computation), sub component fitness evaluation methods, and a “to string” reporting method.

### Main Class

The main driver simply imports and stores the JSON configuration files. It instantiates the EA class. Finally, it initiates the simulation.

### Calculating Fitness

The fitness function for an individual TA pair is defined as the sum of its binding score, nuclease score, deubiquitylating score, NLS score, and T4SS signal score. Each sub-component fitness can maximally be 1 and therefore the max fitness of a perfect TA is 5. To elaborate precisely on how sequence similarity is used as a proxy for fitness, we describe the situation for DUB fitness. The DUB domain is a catalytic sequence of amino acids that conforms to a schema. The DUB schema in *cifs* is precisely, “HWVTLVI---------YY-DSL--------I---L-----D---------QQ-DG---CG----EN”, where dashes (-) are interchangeable spaces (*don’t cares*) and letters are requirements of specific amino acids in specific positions. A perfect alignment score of 1 for a *cif* DUB would match this schema. Anything not conforming to the schema is penalized by the alignment algorithm for gaps and mismatches. The schema for the nuclease domain is as follows, “DL-LL-R----------PIIIELK---------------------DLVL----------PIGLELK”. These two consensus schemas were originally derived directly from compilation of diverse CI and CI-like toxins.^50^ Schemas for NLS and T4SS signals are also pulled from literature and listed on the preceding TA Class description. Thus, by using sequence similarity to conserved schemas and the binding algorithm (described above) we can sum elements for a perfect TA fitness score of 5. Parsimony pressure is applied if a TA genome exceeds a threshold of 4,500 DNA base pairs (this is an estimate of average *cif* TA size) and pressure increases corresponding to the length of the additional extraneous code. Parsimony pressure thus acts to minimize the coding length of TA pairs and accurately reflects selective pressures inducing reduction of *Wolbachia* genomes. In toto, a final fitness score involves the sum of the five functional component scores with a penalty function subtracting a coefficient parsimony penalty based on sequence length. All code is publicly available for inspection and reuse on [github](https://github.com/stampedepress/CytoplasmicIncompatibilityEvolutionaryAlgorithm).

### Experimental setup

The EA evolves populations of TAs and evaluates their fitness. Simulations were initialized via three distinct methods described in **Fig 1C**. How the simulation is initiated impacts the levels of inherent diversity in the starter population. Methods i-iii decrease in starting diversity from most to least respectively. After 1000 generations the simulation is terminated, and data collected. Data collected is given above and was graphed in Graphpad’s Prism software. Experiments were conducted with 30 runs each.

### Statistical Analysis

For experiments generating multiple comparisons like optimizing the EA (**Fig 2** and **3**) we employed one-way ANOVA with Tukey post-hoc analysis using Graphpad Prism software. We compared values present at the final generation at termination of the simulation. P-values were considered significant if less than the standard 0.05. In **Fig 4**. terminal data were compared using unpaired two-tailed t-test with Welch’s correction.

## Results and Discussion

### Validating and Tuning the EA’s Fitness Function

One prerequisite of implementing an EA is an ability to evaluate fitness of individuals. A protein’s function and thus its fitness is encoded in primary structure (amino acid strings). Protein function can be predicted by comparing strings to others with known function. Therefore, we use sequence similarity to *cif* domains as a proxy of fitness and thereby apply selective pressure. In our EA an individual in-silico *cif* is constituted of the two DNA genes and their translated protein strings. Many individual TA pairs are instantiated within populations. The EA mutates and recombines them exactly as DNA can mutate and recombine. Fitness of individual TA pairs is modelled as a sum of 1) how well a toxin can kill a cell (based on sequence similarity to known killer toxin domains from *cins* and *cids*) and 2) how well the antidote binds its partner toxin (modelled as matching charged residues within cognate TA pairs). Additionally, we add 3) NLS and 4) T4SS signal domains as additional summed components of fitness. We then quantified where NLS and T4SS signals evolved during simulations (in *cifA* or *cifB*) and tracked biased emergence of 2×1 versus TA.

After initial design (**Fig 1**), coding, and parameter optimization (**Fig 2**), we determined that the EA evolved efficiently and observed that population sizes of 5000 individuals with offspring sizes of 100 individuals were optimal because they yielded high fitness, diverse outcomes, and fast computation time (**Fig 2A**). These assumptions have flaws (for example nature isn’t an algorithm that optimizes parameters to speed up evolution; discussed below), but these settings served as a starting point. Next, we tested different algorithmic methodologies for parent selection, recombination, mutation, survivor selection, and a “re-instantiation” method immigrating 10%, 1%, or 0% de-novo individuals (described in methods). Results show means and standard deviation from five trial runs after 100 generations. Our goal was to determine optimal algorithms for maximizing *cif* fitness within simulation time periods. After observing EA behavior, we determined to use a “baseline” configuration of K-tournament selection where K=5 for parent selection. Selected parents are transitioned to mating pool where mating only occurs between individuals within that selective sub-population. Mating of TA parents is implemented with 4-point crossover recombination with a self-adaptive mutation rate to generate offspring TAs. The self-adaptive mutation rate is encoded within an individual’s chromosome and can change if higher or lower mutation rates contribute to better fitness. Offspring TAs are loaded back into the main population (µ+λ) and compete for survival via truncation, which culls the lowest fitness individuals. Subsequent experiments used these conditions unless otherwise specified.

### Tracking Evolution of cif Domains Shows that NLS and T4SS Signals are Quick to Evolve

We performed three large experiments based upon three methods of instantiating populations. Our intention was to determine if starting preconditions biased preferential evolution of NLS or T4SS signals in *cifA* versus *cifB*. Any bias might indicate conditions under which 2×1 or strict TA mechanisms would be the result of that given evolutionary process. In these first tests, the EA successfully evolved and evaluated the fitness of TA modules. In all experiments fitness of individuals gradually increased towards 5 (**Fig 3A, F, K**). This control indicates that our code performed as designed. The initial lag observed in **Fig 3K** is a direct result of the instantiation method used, wherein the population was instantiated by mutation from an initial founder with poor fitness. All three simulations show a start at low average TA fitness which improves as more successful TAs evolve and overtake the population.

We tracked each sub-component of fitness including nuclease, DUB, NLS, and T4SS signal evolution (**B, G, L**). These data also indicate our code works correctly as fitness of each sub-component increases with each generation towards a max score of 1. Importantly these data also indicate the inherent complexity of each sub-component and clearly show that NLS and T4SS signals are relatively quick to evolve in simulations (**Fig 3**; green and orange lines respectively). Binding is of intermediate complexity and arises slower (**Fig 3**; yellow lines). Nuclease and DUB catalytic domains are slow to evolve and do not completely reach perfect consensus sequences within the timeline of the evolutionary experiment (**Fig 3**; black and purple lines respectively). These data are in concordance with the given complexity of the domains. For example, the NLS is only 4 residues (“KRAR”) whereas max binding fitness requires 11 matching residues in both toxin and antidote, and consensus sequences of catalytic domains must match 23 conserved residues within their schemas.

The five components’ relative evolvability (or inherent speed of their evolution) indicates that CI systems might frequently lose, replace, adapt, and move NLS and T4SS signals, whereas binding and catalytic domains are more likely to remain conserved in-place due to difficulty of evolving them in the first place. If they are destroyed, they cannot quickly be replaced, whereas NLS and T4SS signals might be “fungible”. We note that a full spectrum of T4SS signals has yet to be identified and these diverse domains perhaps encode room for self-adaptive ambiguity.^52-56^

These data indicate that both strict TA and 2×1 systems could co-exist and might even inter-convert between mechanisms on evolutionary time scales with drift, mutation, and recombination. These in-silico observations are congruent with empirical literature demonstrating both systems are apparently extant.^19,26,29,32,36,37^ Cautiously, we note that these observations are premised on assumptions that there must be some conditions selecting for the evolution of CI; these ecological conditions are not yet completely defined^17,24,25^ yet must exist under some context that gives rise to CI and *cifs;* perhaps amongst discrete spatial limitations and genomic competitions.^2^ Importantly, our model simply justifies how multiple CI mechanisms might evolve to coexist on the amino acid level.

### Parameters of Simulations Bias Evolution of TA versus 2×1

When we measured where NLS and T4SS signals evolved (in *cifA* or *cifB*) under three different starting conditions (random, consensus, and founder; see **Fig 1C** and **D**) we detected biases in the evolutionary trajectory of one model over another (**Fig 3D, I, N, E, J, O**). After random instantiation (method i.) both NLS and T4SS signals’ scores were slightly less than 0.5 indicating a slight preference for evolution of those sequences in *cifA* genes. After semi-random instantiation (method ii.) there was strong bias to evolve the T4SS within *cifA* genes indicating a bias towards 2×1. Only in the third method did both NLS and T4SS signals preferentially evolve in the *cifB* gene, thereby indicating bias towards TA mechanisms. Each method showed statistically different termination conditions for T4SS locations with all p-values < 0.05. Method i significantly differed from method iii with respect to termination condition of NLS signal. Importantly, these results indicate that our model can detect significant evolutionary bias towards one mechanism over another.

We next sought to understand the conditions that drove biased evolution of one mechanism over another. To monitor genetic diversity within the populations we tracked a diversity index, which was determined by randomly sampling 10 toxins from the population each generation and calculating the average similarity of those ten toxins’ amino acid strings. In populations where individual TAs fix and overtake the population, diversity decreases to zero (**Fig 3C, H**, and **M**). After 1000 generations, most populations are overtaken by one or a few TAs of high fitness. In method iii, which resulted in biased evolution of strict TA systems, the diversity index was lowest (see **Fig 3M**). We tested whether diversity directly drove bias by altering the relative levels of genetic diversity within the population. We controlled this by simply changing population size (µ). Smaller populations carried less diversity (**Fig 4A**). The relative diversity did alter the course and path of evolution, but not the outcome, which converged (**Fig 4B** and **C**). We next tested whether length of the toxin protein impacted the outcome. When we doubled the size of the average size of the toxins, we significantly raised the bias of the model towards the TA model (**Fig 4**). We discuss the theoretical impacts of these observations below.

Importantly, these observations demonstrate that we have successfully encoded an EA that evolves and tracks *cif* amino acid evolution.

Overall, our model sheds some light about conditions that might bias the evolution of one model over another and hence explain why four studied orthologs appear to operate as strict TAs, but two orthologs seemingly behave in a 2×1 fashion. Only the third instantiation method exhibited tandem bias of NLS and T4SS evolution in *cifB* genes, indicating strong preference for a strict TA functionality. Notably, this model had the least diversity within its population and likely reflects more accurately the actual evolution in *Wolbachia* systems where an insect is colonized by a founder strain and diversity is only rarely encountered in sporadic co-infections that only occur rarely in evolutionary history, but are likely the source for CI gene evolution if phages exchange genes during coinfection. Therefore, our analysis can explain the observed bias in favor of strict TA functionality by about 60% of studied *cif* orthologs; notably in our simulation method iii) the bias was also about 60%. To be cautious, however, we note that only ∼6 ortholog TA pairs have been studied in detail and it remains to be seen whether the observed frequency of TA or 2×1 functionalities is some relic of sample bias. Future studies will utilize this framework to determine more conditions that give rise to 2×1 versus TA systems.

To elaborate on the question of which instantiation method is more biologically realistic, we suggest the following thoughts. One assumption is that a prokaryotic TA model preceded the evolution of CI. There is evidence for this in the fact that *cif* operons have been observed within plasmids of *Rickettsia* (a sister lineage of *Wolbachia*), lending plausibility to the hypothesis that CI emerged from an ancestral prokaryotic TA plasmid selection system.^50^ If this hypothesis is correct, method i) random instantiation, is not biologically relevant as it begins selection on all four components simultaneously from completely random sequences. In contrast, method ii) assumes a prokaryotic TA system already exists and reasonably comports with some known toxin consensus sequences then selects for the addition of NLS and T4SS signals in the jump from prokaryotic TA to eukaryotic CI. This method is biologically relevant only for the first emergence of CI’s evolution in deep evolutionary history. In that situation and according to the data observed here, this model shows preference for the evolution of 2×1 systems, but not to the complete exclusion of TA systems. After emergence of TAs, our model’s data predicts they flux periodically from 2×1 to strict TA.

### Increasing Length of Toxin Biases Evolution Towards TA mechanisms

Instantiation method iii), where a population is generated by a founder, more accurately reflect the day-to-day evolution of *Wolbachia* organisms in their hosts. In each insect, the *Wolbachia* encountered will be entirely derived from the ancestor of that infection and therefore recombination with sequences of radically different *cifs* is unlikely, though not impossible due to mobility of WO phage viruses and infrequent co-infections. Method iii) most accurately reflects these conditions and in this model, there was strong bias towards the evolution of strict TA functionality (**Fig 3**). This suggests that over time, most (∼3/4) CI systems should end up in a state of strict TA functionality with some variation induced by ongoing flux of NLS and T4SS signals. One of the key factors seemingly controlling this evolution is simply the length of the corresponding antidote and toxin (**Fig 4**). Because the NLS and T4SS signals are of low complexity and evolve quickly, stochastically they should simply arise more often in what is the longer gene of the pair. Of all syntenic *cif* operons, the length of the toxin is always longer than the length of the antidote. This also indicates a simple bias toward strict TA if NLS and T4SS signals simply drift into the larger ORF. Biology is complex, yet factors having the biggest role in these mechanistic biases might be as simple as gene length. However, sequence length doesn’t explain everything about the model. In data from method ii), where the strongest bias towards 2×1 was observed, the toxins and antidotes on average are the same size. Therefore, size does not account for all the forces driving bias in either direction.

### Conclusions, Future Directions, and Limitations

The hypotheses and take-homes from our model are thus: 1) CI might evolve from less complex prokaryotic TA systems (**Fig 1**). 2) TA systems can convert to CI systems by the addition of at minimum NLS and T4SS signals (Fig 1) though these domains may not be completely sufficient. 3) Where NLS and T4SS signals evolve (in cifA or cifB) is predicted to be the determinant of 2×1 or strict TA mechanics (Fig 1). 4) In cases where CI evolves from a pre-existing TA framework, the evolution is biased towards 2×1 systems, but not at full exclusion of strict TA (Fig 3I and J). 5) In our model, sequence diversity impacts the path of evolution, but not the terminal result, whereas sequence length can predispose bias of signal evolution in a location. Finally, 6) Codon-based EAs can be applied in a bottom-up approach to address questions related to the evolution of protein strings.

In future experiments we plan to utilize this framework to test additional sequences of NLS and T4SS signals. Importantly, the signals we used are not the only ones that exist in nature. There can be cryptic and/or bi-partite combinatorial sequence motifs that contribute to secretion and localization.^55^ To add to the nuance, our algorithm doesn’t account for redundant sequences. For example, it doesn’t quantify if additional NLS or T4SS signals evolve elsewhere, beyond the first. It would be interesting to re-program the system to measure and tally if multiple NLS and T4SS signals are evolving and where they are. One prediction our model makes is that because NLS and T4SS signals are of low complexity, there may be multiple redundant signals within the same gene. In future experiments we will look for this.

Our model makes many assumptions. One assumption we made, to begin analysis somewhere, is that the parameters causing the fastest evolution of *cifs* in simulations were apt to simulate the natural evolutionary dynamics of these TA modules. However, evolution within the natural organism might not be so ideal.

Therefore, favoring the most efficient methodologies and parameters to evolve high fitness quickly might be incongruent with nature. Although we grounded the evolution of the EA in real biology in the code using actual *cif* sequences and known binding features. The benefit of our coded framework is that it can be modified to test and address future criticisms and hypotheses. For example, while we’ve only implemented the 2×1 and strict TA mechanisms, if ever a possibility of a third mechanism is observed or postulated, we can add that possibility to the code base.

Finally, our model encodes and models evolution of the most primal or basal level of CI (the amino acids). It is not an ecological model assessing TA allele fixation in populations. It would be inappropriate to directly compare our model with prior ecological models;^2,17^ although our model could be imported into those models as an foundation. The natural evolution of selfish TA elements involves multiple levels of evolutionary dynamics. For example, *cif* systems exist within WO-phages that exist within *Wolbachia* bacteria that live within insect hosts that live within populations. *Cifs* impact evolution and population dynamics on all these levels. Future models might incorporate our codon-based EA as a subcomponent of a larger multi-competitive EA framework. Such a program might provide vast insights into the complex evolutionary dynamics inherent to *Wolbachia* biology and make predictions about actual CI gene function.

## Acknowledgments

We thank Jason Barbieri, David Edwards, and Dennis Brown for constructive comments on the project design and manuscript. Funding was provided by Auburn University’s Department of Entomology and Plant Pathology startup funds (JFB) and US Department of Agriculture Hatch Grant (1015922). JJG acknowledges support from the NIH/NIAID (R21 AI146773, R21 AI156762, R21 AI166832).

## Data Availability Statement

All code and datasets are available on [Github](https://github.com/stampedepress/CytoplasmicIncompatibilityEvolutionaryAlgorithm).

## Conflict of Interest

Authors have no conflicts of interest to report.

## Footnotes

1 https://www.python.org/

2 https://biopython.org/

## Notes

### Competing Interest Statement

The authors have declared no competing interest.

